# Genome Skimming with Nanopore Sequencing Precisely Determines Global and Transposon DNA Methylation in Vertebrates

**DOI:** 10.1101/2023.01.25.525540

**Authors:** Christopher Faulk

**Author notes:** Denotes corresponding author. Contact.

## Abstract

Genome skimming is defined as low-pass sequencing below 0.05X coverage and is typically used for mitochondrial genome recovery and species identification. Long read nanopore sequencers enable simultaneous reading of both DNA sequence and methylation and can multiplex samples for low-cost genome skimming. Here I present nanopore sequencing as a highly precise platform for global DNA methylation and transposon assessment. At coverage of just 0.001X, or 30 Mb of reads, accuracy is sub-1%. Biological and technical replicates validate high precision. Skimming 40 vertebrate species reveals conserved patterns of global methylation consistent with whole genome bisulfite sequencing and an average mapping rate above 97%. Genome size directly correlates to global DNA methylation, explaining 44% of its variance. Accurate SINE and LINE transposon methylation in both mouse and primates can be obtained with just 0.0001X coverage, or 3 Mb of reads. Sample multiplexing, field portability, and the low price of this instrument combine to make genome skimming for DNA methylation an accessible method for epigenetic assessment from ecology to epidemiology, and by low resource groups.

## Introduction

Genome skimming refers to unbiased low-pass sequencing below 0.05X coverage and is used to reconstruct mitochondrial genomes as well as for species and parasite identification.^1–3^ Oxford Nanopore Technologies’ nanopore sequencers have been used for genome skimming to produce mitochondrial genomes with success, however no studies have used genome skimming to report DNA methylation.^4^ An important biomarker, DNA methylation varies with age, tissue, species, and environmental exposures making it a useful measure from ecology to epidemiology.^5^

Global DNA methylation measures the ratio of 5’methylcytosines (5mC) vs. total cytosines reported as a percentage. Current methods to measure global methylation have drawbacks in either cost or accuracy. Antibody based methods do not have the resolution to distinguish small magnitude changes often seen in biologically significant epidemiological exposures.^6^ Other methods such as reduced representation bisulfite sequencing (RRBS) and whole genome bisulfite sequencing (WGBS), while accurate, are expensive, biased in target location, and rely upon bisulfite conversion which causes downstream challenges. Bisulfite conversion chemically modifies unmethylated cytosines to uracils, read by polymerases as thymines. In the reaction DNA is sheared to very short fragment length. During analysis, alignment becomes challenging due to the shift from a 4-base alphabet to a mainly 3-base encoding, necessitating special aligners and loss of the ability to detect mutations in the native genome sequence.

Nanopore sequencers produce long reads from native genomic DNA, typically in the 10-50 Kb range, and simultaneously report the presence of DNA methylation genome-wide.^7^ Long reads enable more contiguous genome assembly and easier reconstruction of repetitive regions.^8^ Alignment is faster with fewer errors with the ability to span large transposons and detect structural variants and single nucleotide variants. Applied to genome skims, long reads enable more accurate recovery of mitochondrial genomes with lower sequencing depth.^9^ Sample multiplexing, field portability, and the low price of the instrument all reduce cost and enable sequencing by low resource groups.^10^

Genome skimming is unbiased in most genomic regions; however, some regions are present in multiple copies and this enrichment will be reflected in the skimmed read dataset. Mitochondria can be present in thousands of copies per nuclear genome and are therefore a common analysis target since a skimming level of even 0.05X would result in hundreds of full-length copies of mitogenomes. Transposons are another overrepresented category in the genome. Repeats make up ∼50% of a mammalian genome on average, with some families present in over a million copies per nuclear genome.^11^ Though few studies have focused on transposons in genome skimming data, the potential is high for transposon biology.^12–14^ From an epigenetic perspective, transposons have long been used as proxies for global DNA methylation in epidemiology, so validation of their methylation to global DNA methylation holds promise for their continued use as biomarkers with a lower cost per sample.^15^

Here I present nanopore sequencing as a highly precise, widely applicable platform for global DNA and transposon methylation assessment. Coverage depth proves its accuracy down to skimming levels of 0.001X, or 30 Mb in mammalian genomes. Both biological and technical replication validates measurement precision across tissues and by low and high methylation controls. Comparative genome skimming from 40 different species across the vertebrate radiation reveals conserved patterns of methylation and serves as evidence for wide applicability. Transposon methylation is accurately and repeatedly determined in both mouse and primates with major classes showing precision at just 0.0001X coverage, or 3 Mb of sequencing.

This is the first report of nuclear DNA methylation assessment using genome skimming. Nanopore sequencing is revitalizing genome skimming in ecology with detection of cryptic hybridization of threatened primates and in non-invasive methods to assemble mitogenomes^16,17^. Extending its utility by adding DNA methylation capability will aid the use of genome skimming in epidemiology, conservation, and comparative genomics.

## Results

### Coverage Depth Estimation

Prior to designing a genome skimming experiment it is necessary to determine the minimum level of sequencing depth to obtain sufficient precision for statistical power. A single chimpanzee genome was sequenced to 11.2X depth using a nanopore PromethION instrument and global DNA methylation assessed at 77.87%. Reads were subsampled at coverage levels representing the range of genome skimming expectations, from 0.1X (300 Mb) down to 0.0001X (300 Kb) and bootstrapped 10 times to calculate error (Table 1). At 0.01X (30 Mb), coverage of a primate genome results in an error of less than 1% difference from the true value (Figure 1).

**Table 1.** DNA Methylation at Varying Coverage Levels, Bootstrapped Subsampling

|  | 11.2X<br>(34 Gb) | 10X<br>(30 Gb) | 1X<br>(3 Gb) | 0.1X<br>(300 Mb) | 0.01X<br>(30 Mb) | 0.03X<br>(10 Mb) | 0.001X<br>(3 Mb) | 0.0001X<br>(300 Kb) |
| --- | --- | --- | --- | --- | --- | --- | --- | --- |
| Mean | 77.87 | 77.89 | 77.91 | 77.98 | 77.88 | 77.92 | 77.79 | 78.6 |
| Bootstraps | 1 | 10 | 10 | 10 | 10 | 10 | 10 | 10 |
| Minimum |  | 77.88 | 77.8 | 77.76 | 77.25 | 76.51 | 75.08 | 69.44 |
| Maximum |  | 77.9 | 77.97 | 78.21 | 78.42 | 79.49 | 81.8 | 88.22 |
| Range |  | 0.0182 | 0.168 | 0.4527 | 1.17 | 2.977 | 6.72 | 18.78 |
| Std. Deviation |  | 0.0061 | 0.046 | 0.1757 | 0.4176 | 1.053 | 2.187 | 6.241 |
| Std. Error |  | 0.0019 | 0.015 | 0.05556 | 0.132 | 0.3329 | 0.6914 | 1.973 |

**Figure 1:**
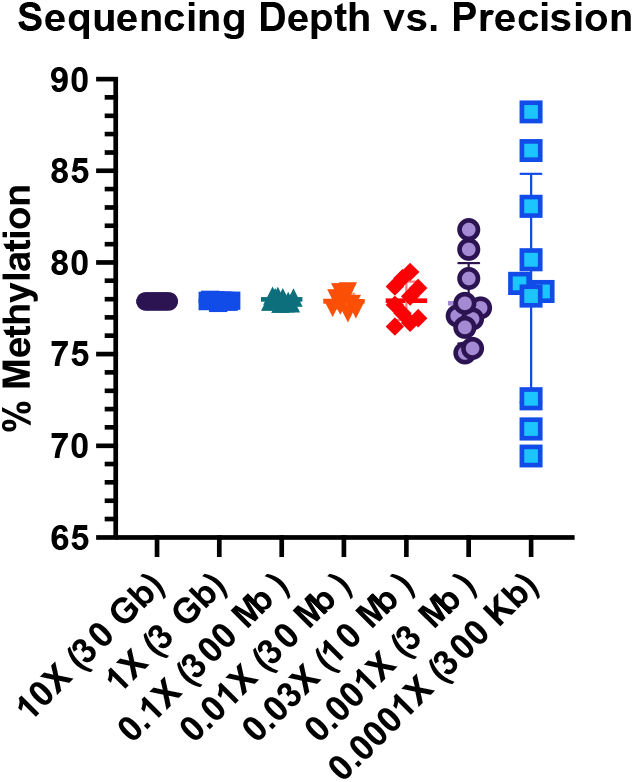
Sequencing depth vs. precision. Reads are subsampled from an 11.2X coverage chimpanzee genome at coverage levels from 0.0001X to 10X. DNA methylation is calculated for 10 bootstrapped subsamples.

### Biological and Technical Replication

Reproducibility is crucial for assays which may vary over a small range, and is measured by precision, i.e., how close repeated measures are to each other. In the following experiment, mice were sequenced to skimming level coverage between 0.0025X - 0.016X (median 0.0058X) or by read coverage, from 8 Mb - 133 Mb (median 47 Mb) reads per sample (Table 2). Three tissues per mouse were sequenced with five mice serving as biological replicates. For controls, whole genome amplified DNA is expected to have near zero DNA methylation and was used as the low methylated control, while CpG methylase treated DNA is expected to have near 100% methylation and was used as the high methylated control. Multiple aliquots of the same two control reactions were individually barcoded along with the mouse tissues and run on a single MinION flow cell. These served as technical replicates as well as range controls.

**Table 2.** Biological and Technical Replication of DNA Methylation in Mouse at Low Coverage.

| Tissue | Methylation (%) | SD (%) | Replicates | Median (Mb) | Mean % Mapped | Mean Quality |
| --- | --- | --- | --- | --- | --- | --- |
| Hippocampus | 72.18 | 1.24 | 5 | 28.3 | 98.81 | 30 |
| Cortex | 72.75 | 0.80 | 5 | 13.5 | 98.97 | 29.1 |
| Testes | 75.30 | 0.86 | 5 | 48.5 | 98.83 | 30.2 |
| Low Control | 1.48 | 0.13 | 5 | 99.2 | 99.18 | 26 |
| High Control | 94.82 | 0.19 | 4 | 51.0 | 98.96 | 30.1 |

DNA methylation was highly consistent within tissues and significantly higher in testes vs. hippocampus and cortex (p<0.001) (Figure 2). Quality control validated the experimental protocol (Figure 3). Methylation values were robust to differences in total bases and average length sequenced, with a 4-fold difference in total reads from the lowest group to the highest, and a 2-fold difference in read length between groups. Quality scores did not affect methylation accuracy either, with cortex having a significantly lower, but still high-quality average q-score between tissues (29.1 vs. 30.2). The lowest q-score occurred in the low methylation control, likely due to the poorer quality of the *in vitro* whole genome amplified DNA. Both controls performed as expected, though the high control was short of 100% due to incomplete enzymatic conversion; its technical replication proved very precise with a 0.19% standard deviation. All reads reliably mapped to the mouse genome with less than 0.37% difference across all 24 samples. Mean read quality was slightly lower for cortex than hippocampus or testes but did not affect precision.

**Figure 2:**
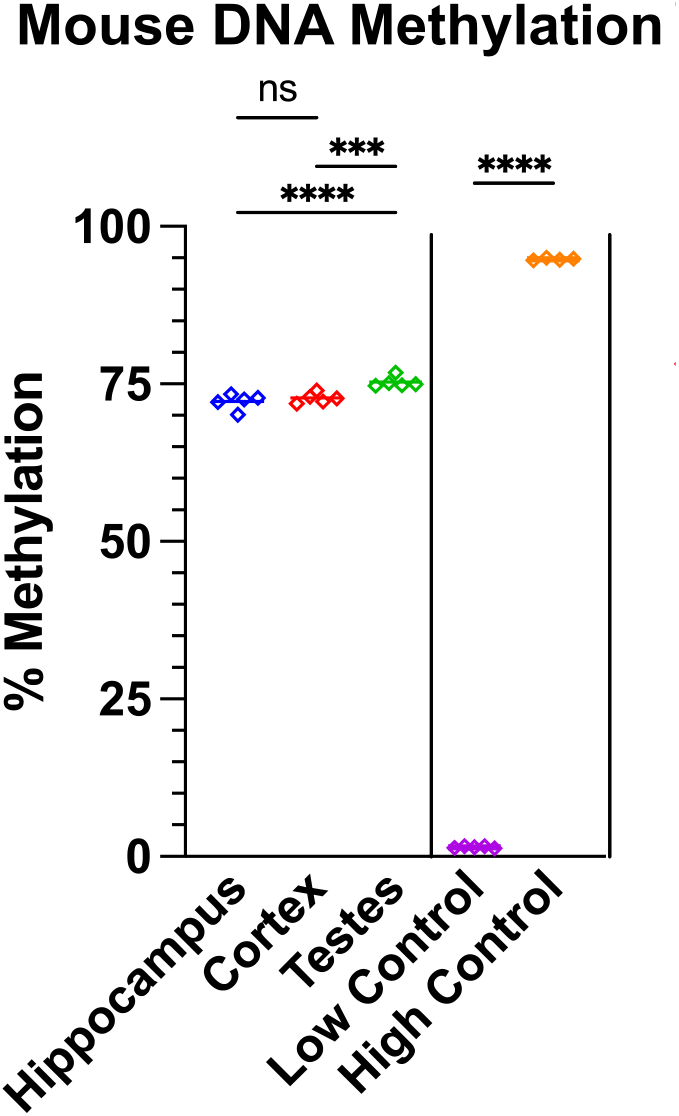
Replicate measures of DNA methylation in mouse. Hippocampus, cortex, and testes are biological replicates. Control samples are technical replicates. Significance is compared within tissues and controls but not between them (*p ≤ 0.05, ** p ≤ 0.01, *** p ≤ 0.001, **** p ≤ 0.0001).

**Figure 3:**
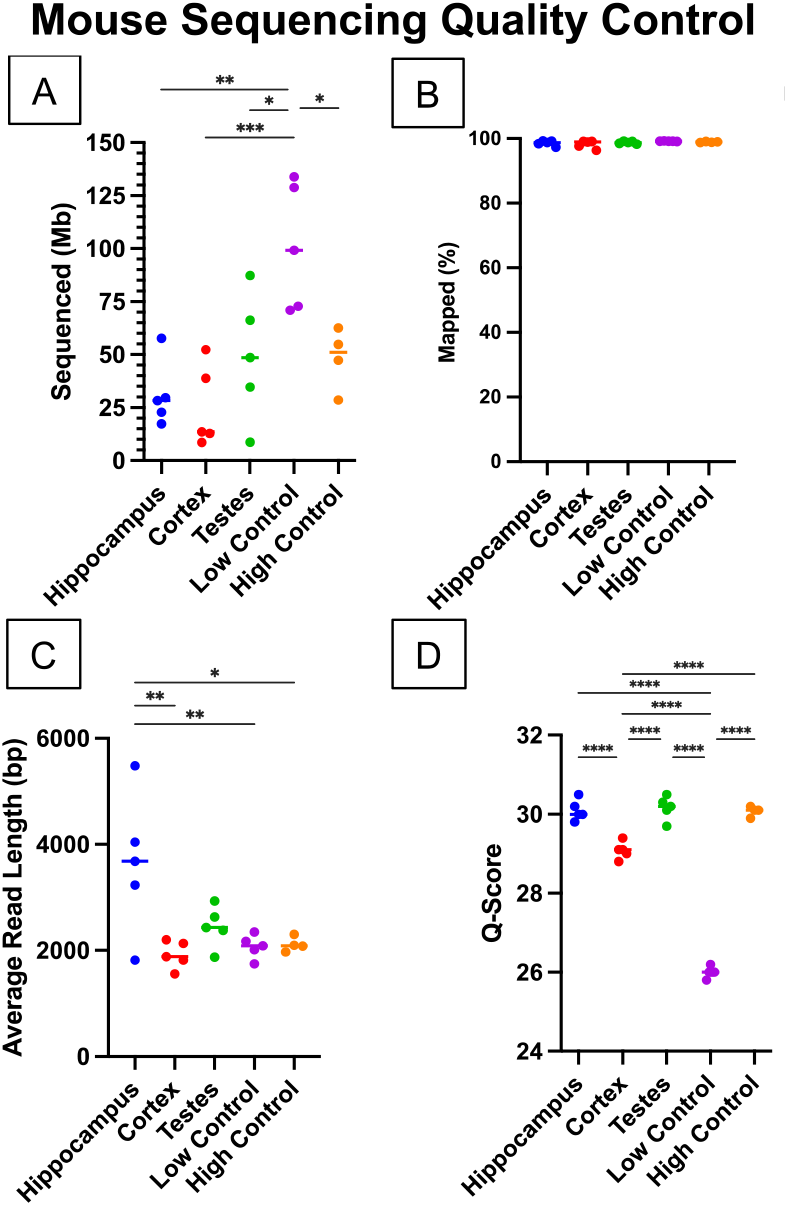
Mouse sequencing quality control. (A) Number bases sequenced in Mb. (B) Percentage of bases mapped to mm39 reference genome. (C) Average read length in base pairs. (D) Average read quality scores (*p ≤ 0.05, ** p ≤ 0.01, *** p ≤ 0.001, **** p ≤ 0.0001).

### Vertebrate methylation

Methylation varies with tissue and by species. To determine whether skimming is sufficient to capture primate-specific levels of global DNA methylation and recapitulate phylogeny, I sequenced buffy coat DNA from five primates to high depth and downsampled the reads to 0.01X equivalent, or 30 Mb (Figure 4). The global patterns match genetic distance from chimpanzee, though the two macaque species differ from each other, despite roughly equal time of divergence from their common ancestor with humans.

**Figure 4:**
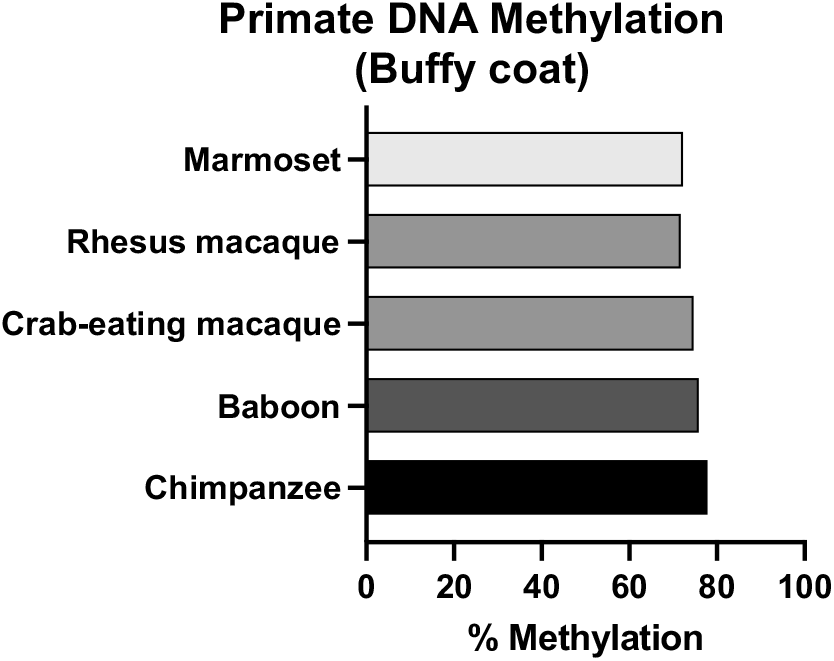
DNA methylation from primate buffy coat. The bars are colored by genetic distance from human, dark to light.

Across vertebrates, methylation varies dramatically. Muscle tissue from 34 species was sequenced at depth ranging from 3 to 200 Mb, median 80 Mb (Figure 5a). Genome size correlated with percent methylation with an R^2^ of 0.44, likely due to increased repetitive element content and associated silencing. (Figure 5b). Summary statistics are shown in Table 3, complete data are available in supplementary file 1.

**Table 3.**
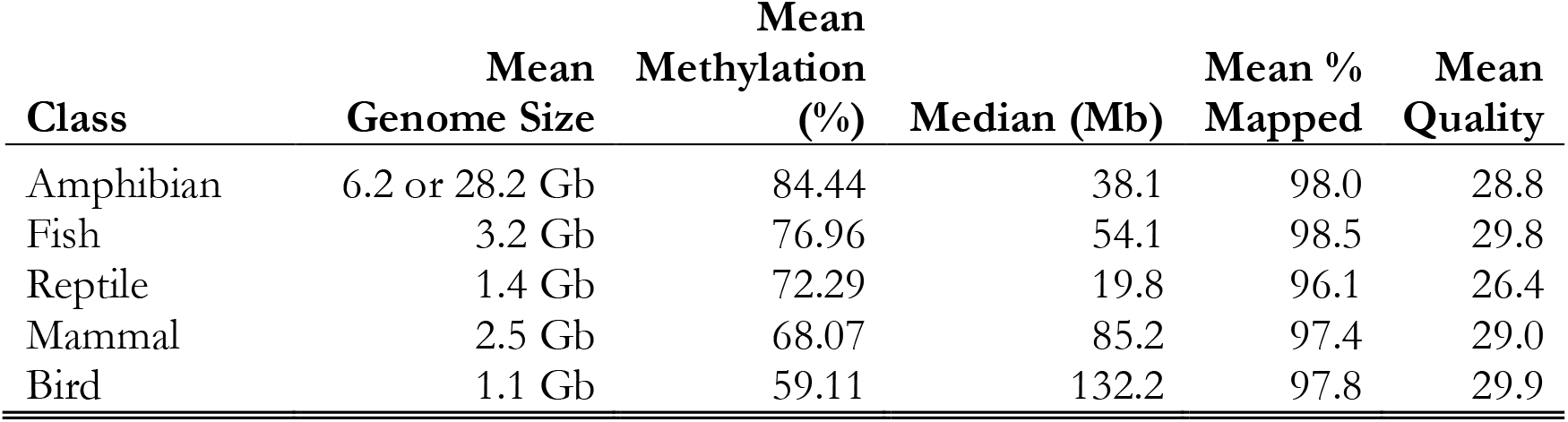
Summary Statistics for Vertebrates by Class

| Class | Genome Size | Mean Methylation (%) | Median (Mb) | Mean % Mapped | Mean Quality |
| --- | --- | --- | --- | --- | --- |
| Amphibian | 6.2 or 28.2 Gb | 84.44 | 38.1 | 98.0 | 28.8 |
| Fish | 3.2 Gb | 76.96 | 54.1 | 98.5 | 29.8 |
| Reptile | 1.4 Gb | 72.29 | 19.8 | 96.1 | 26.4 |
| Mammal | 2.5 Gb | 68.07 | 85.2 | 97.4 | 29.0 |
| Bird | 1.1 Gb | 59.11 | 132.2 | 97.8 | 29.9 |

**Figure 5:**
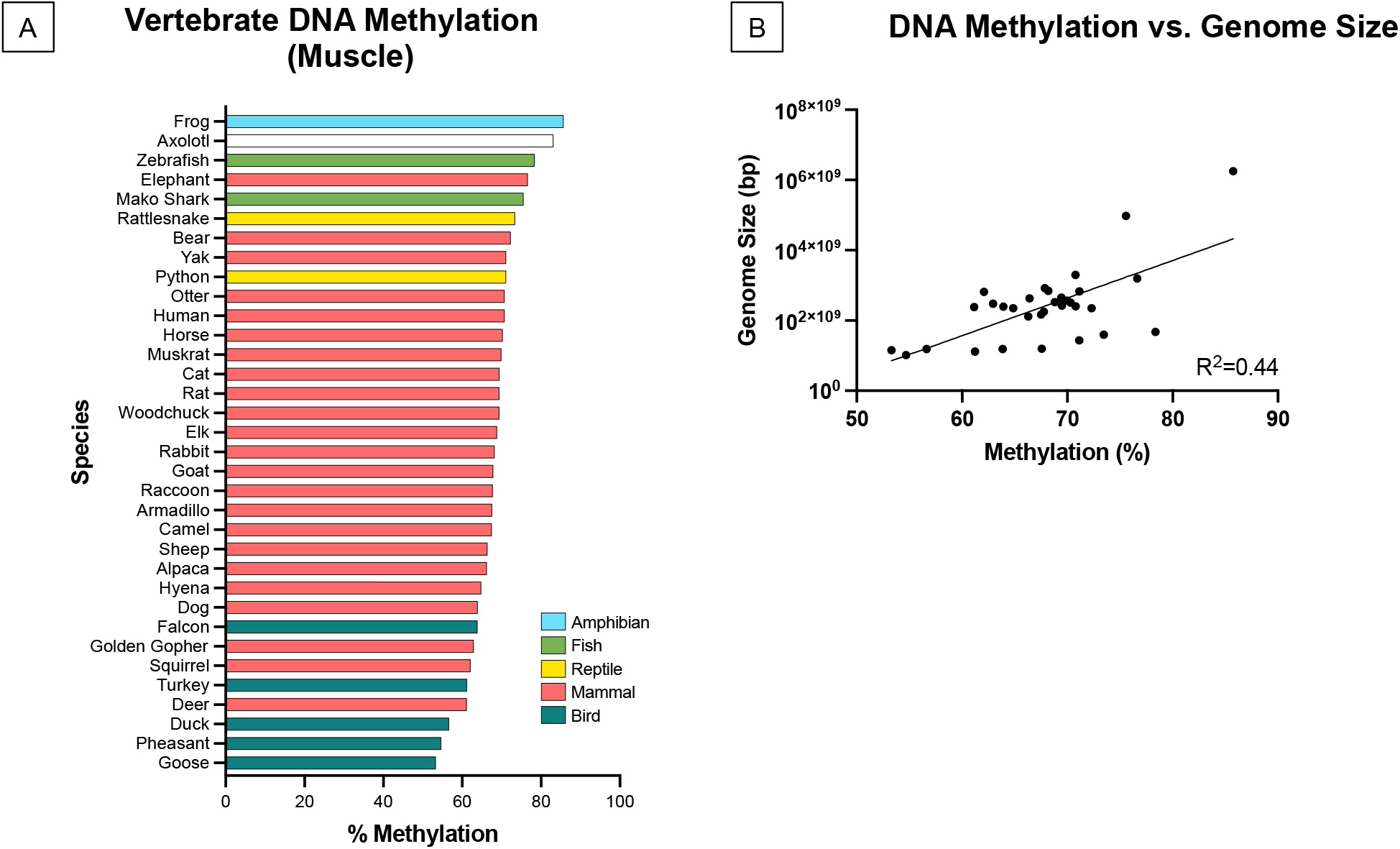
Vertebrate DNA methylation. (A) Global DNA methylation from skeletal muscle in vertebrates. (B) Genome methylation percentage vs. genome size in base pairs. Axolotl was removed as an outlier, as it is an order of magnitude larger than other genomes.

### Transposons

Genome skimming is unbiased by nature; however, some regions of DNA are enriched compared to single copy genes, for example, mitochondria which are the traditional target of genome skimming. Repetitive elements and transposons are present in multiple copies per cell and therefore are more likely to be sequenced by skims than single copy regions. Importantly, DNA methylation plays a strong role in suppressing transposon mobilization and genomic dysregulation. I reasoned that with ∼40% of a typical mammalian genome consisting of repetitive elements, genome skimming can assess methylation at the transposon family level by averaging methylation across all copies of a family.

To determine the minimum depth necessary for precise quantitation, I used the high depth chimpanzee dataset downsampled from 10X to 0.0001X, quantified DNA methylation, and bootstrapped 10 replicates (Figure 6 & Table 4). For *Alu* elements, which number over 1 million copies per cell, even 0.001X coverage, just 3 Mb sequenced, is sufficient for sub-1% accuracy. LINE1 elements are present in 100,000 copies per cell, though most are truncated. Methylation at LINE1 at the 0.01X coverage level, 30 Mb sequenced, obtained sub-1% precision. Unsurprisingly, *Alu* mean methylation was very high at 92.48%, higher than both LINE1 (84%), and the overall genome (77.87%). These values correlate with transposon age and CpG density, where *Alus* are younger and more CpG dense than LINE1s, and both are under selection for increased DNA methylation to suppress mobilization.

**Table 4.**
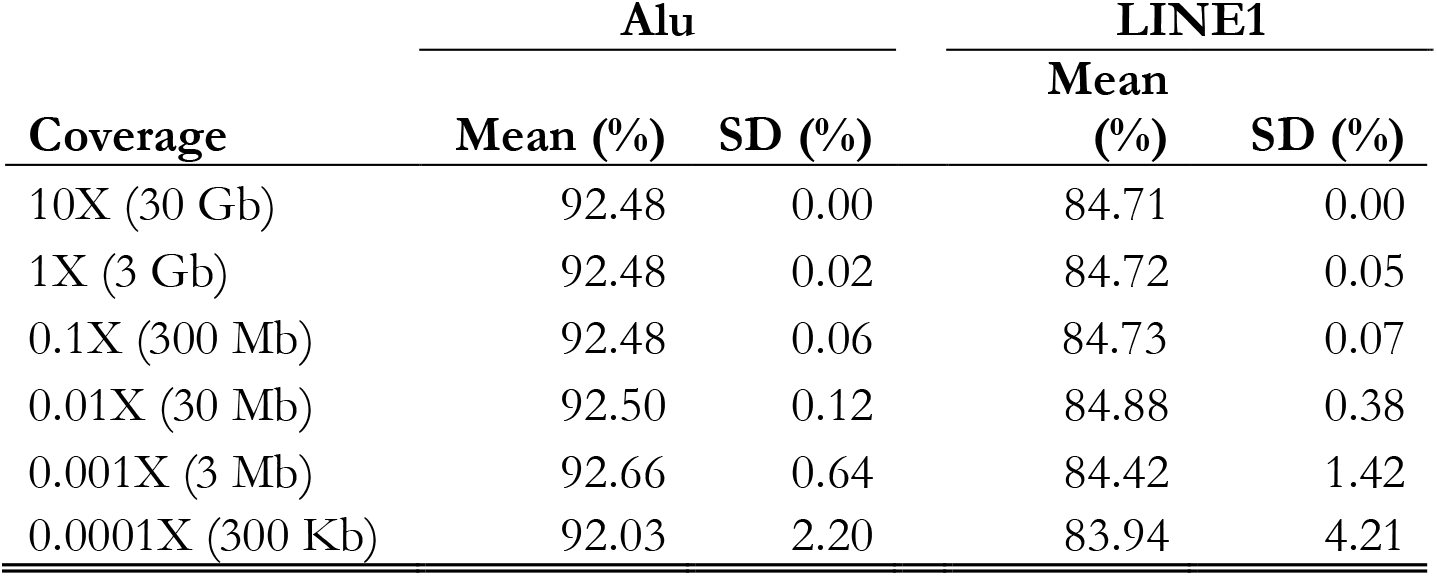
Chimpanzee *Alu* and LINE1 Methylation with 10 Bootstrap Replicates

| Coverage | Alu |  | LINE1 |  |
| --- | --- | --- | --- | --- |
|  | Mean (%) | SD (%) | Mean (%) | SD (%) |
| 10X (30 Gb) | 92.48 | 0.00 | 84.71 | 0.00 |
| 1X (3 Gb) | 92.48 | 0.02 | 84.72 | 0.05 |
| 0.1X (300 Mb) | 92.48 | 0.06 | 84.73 | 0.07 |
| 0.01X (30 Mb) | 92.50 | 0.12 | 84.88 | 0.38 |
| 0.001X (3 Mb) | 92.66 | 0.64 | 84.42 | 1.42 |
| 0.0001X (300 Kb) | 92.03 | 2.20 | 83.94 | 4.21 |

**Figure 6:**
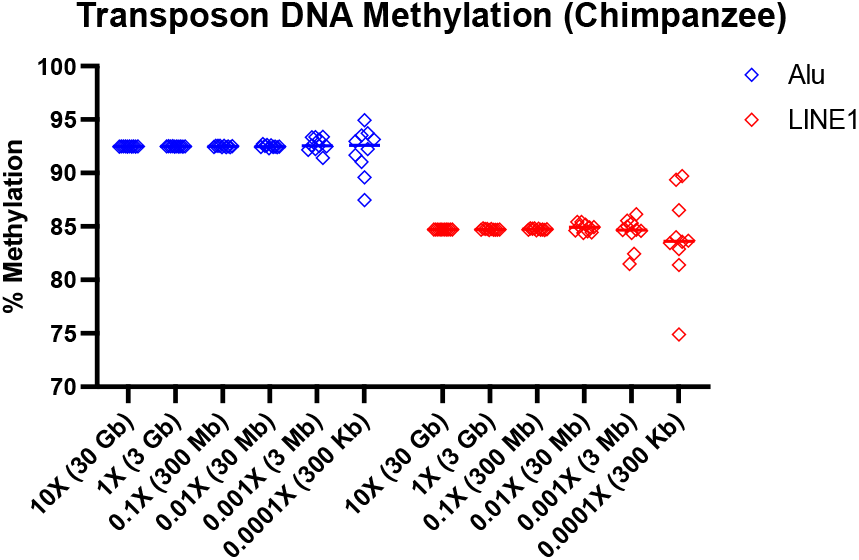
Transposon DNA methylation from chimpanzee buffy coat. From a single 11.72X coverage chimpanzee genome, 10 subsamples of the genome were taken according to coverage level. Alu and LINE1 methylation are shown for the 10 bootstrap replicates.

The mouse genome skims provide validation and biological replication for transposon methylation assessment. I selected the highest copy number families in mouse, with a 2% genomic content threshold and compared their variability within and across tissues (Figure 7 and Table 5). Above the threshold are SINE families, Alu-B1, B2, and B4; ERV families, class II and ERV-MaLR, and LINE family L1. As in primates, all transposons had higher methylation than the global average of 72.18 - 75.30%. Alu-B1 and L1 elements were stable across tissues while B2s, B4s, and ERVL-MaLRs were significantly increased in testes.

**Table 5.**
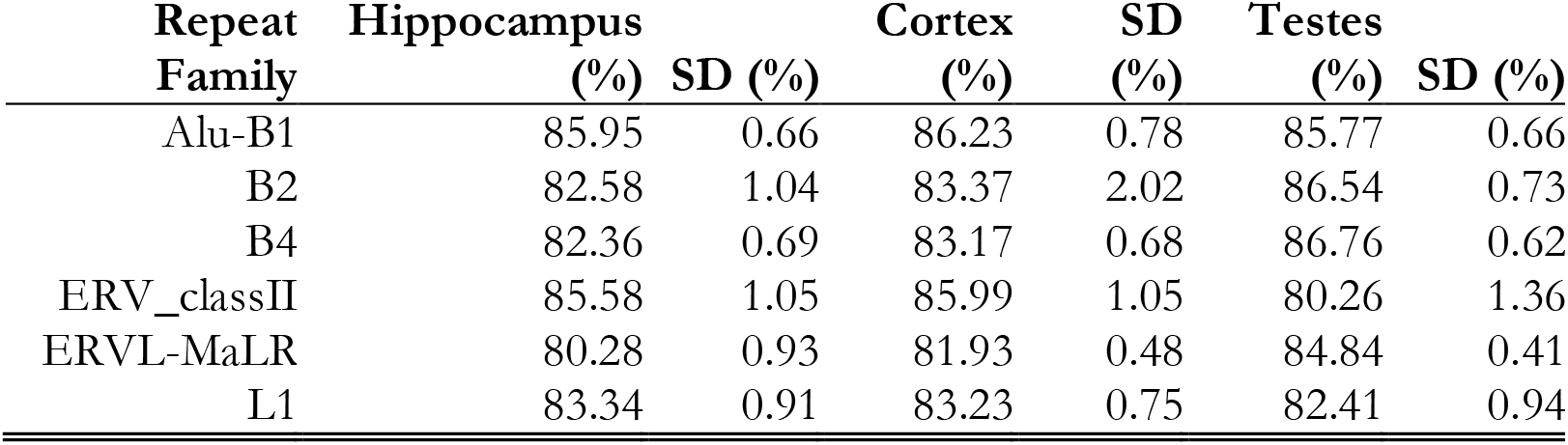
Mouse Repeat DNA Methylation by Family and Tissue

**Figure 7:**
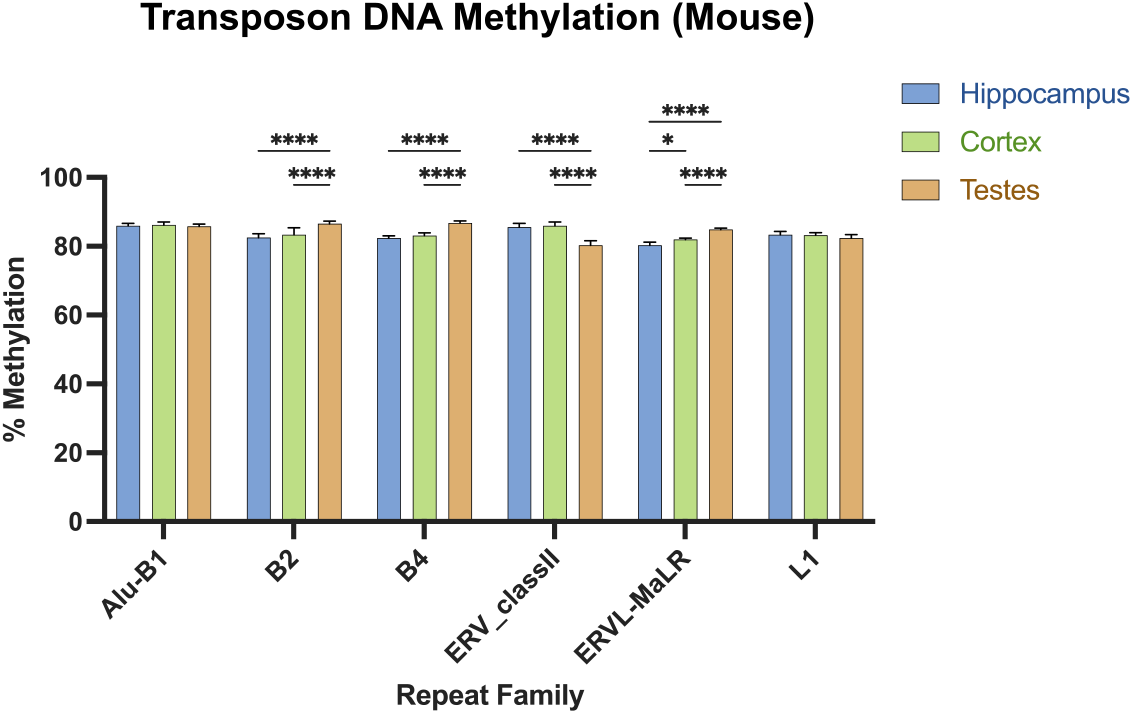
Transposon DNA methylation from mouse. Transposon families above a threshold of 2% genomic content are shown. DNA methylation is calculated from biological replicates of mouse tissues with coverage ranging from 0.0025X to 0.016X. Transposon families with significant differences are shown with ANOVA followed by Tukey’s multiple comparison test. (*p ≤ 0.05, ** p ≤ 0.01, *** p ≤ 0.001, **** p ≤ 0.0001).

ERV_classII elements were the only family with significant decrease in testes, which is biologically significant as the most frequently active transposon in mice is the intracisternal A particle (IAP), an ERV class II element.

By comparing the methylation of all repeat families across species, patterns emerge (Figure 8). The mouse repeat complement shows highest methylation in LTRs, SINEs (Alu-B1, B2, B4), ERVs, and LINE1, which are the most active and prevalent transposons in the mouse genome. Similarly in the chimpanzee, SINE *Alu* and LINE1 are among the most highly methylated and are the most prevalent in the primate genome. Chimpanzee ERVs remain highly methylated despite having lost mobility in the primate lineage. In both mouse and chimpanzee, non-transposon repeats have the lowest levels of methylation, e.g. low complexity repeats, tRNAs and simple repeats. Note that the mouse is represented by 5 replicates taken from hippocampus, while the chimpanzee is represented by buffy coat DNA taken from a single individual.

**Figure 8:**
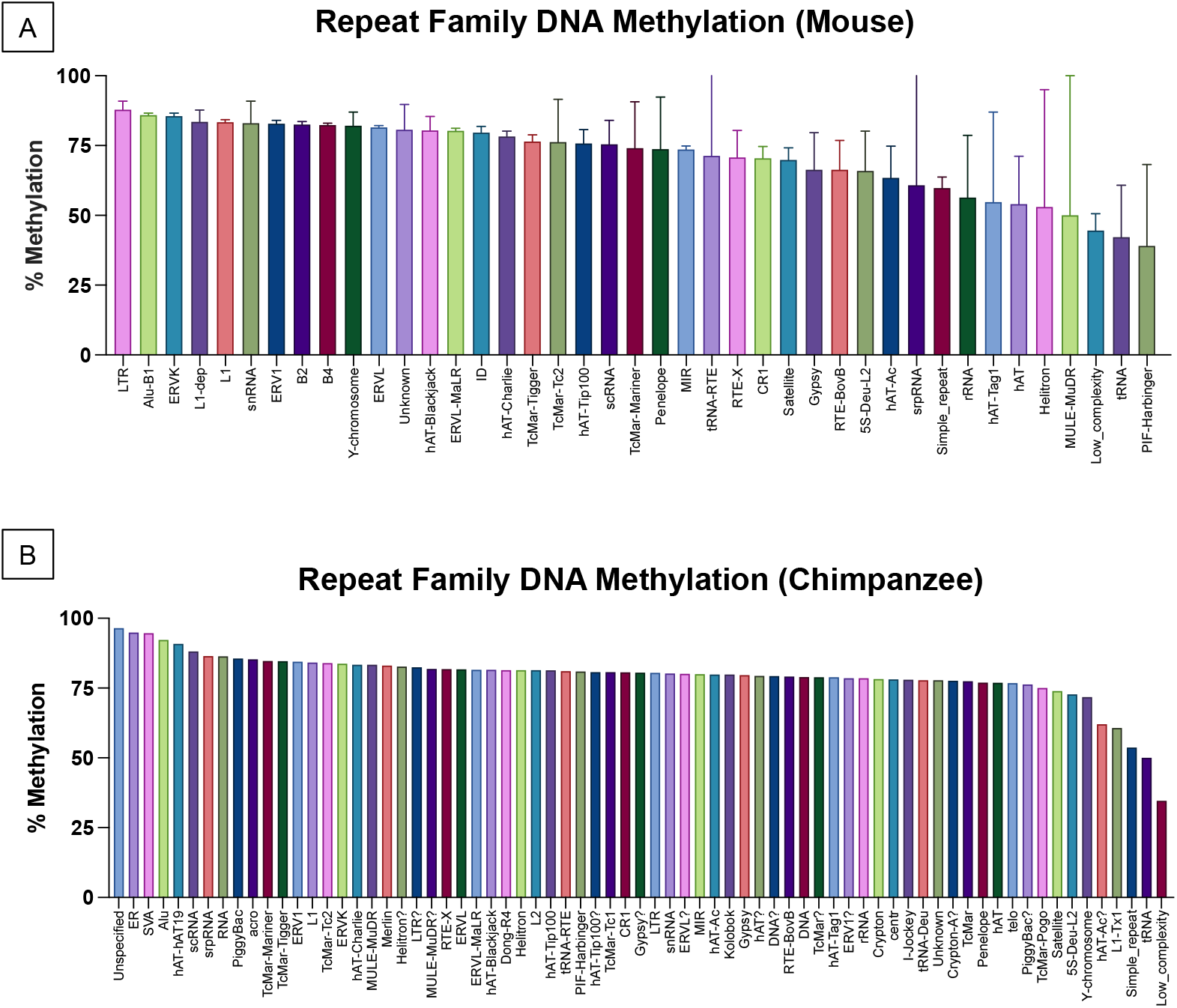
All repeat family DNA methylation. (A) All repeat families from mouse are shown with replication error indicated by error bars. (B) DNA methylation from chimpanzee is shown by repeat family, represented by a single individual.

## Discussion

### Coverage Depth Estimation

Genome skimming, also called low pass or low coverage sequencing, is defined as shallow sequencing down to 0.05X coverage of a genome.^18^ Here I determined that just 0.01X coverage is sufficient to achieve high precision in global DNA methylation assessment in vertebrate genomes. Genome skimming has been used for diverse purposes including plastid genome assembly, parasite identification, evolutionary biology in extinct species, and is gaining software tools for phylogenetic analyses.^3,19–22^ Classical genome skimming focuses on mitochondrial genome recovery for species or individual identification but does not typically utilize the nuclear portion of the skimmed reads or their DNA methylation. There are only five reports of nanopore sequencing used for genome skimming in animals at the time of publication, none of which examine the epigenome.^4,23–26^ In only one case has genome skimming been used to reconstruct mitochondria with a methylation-aware base caller but they did not assess either mitogenome or nuclear DNA methylation.^23^ The generation of long-reads is crucial to improve alignment over traditional short reads.^27^ The further use of nanopore sequencers for genome skimming will enable the acquisition of DNA methylation in parallel to the DNA sequence data for no additional cost.

Genome skimming for DNA methylation is applicable to any method that generates nanopore sequence data even in small quantities such as field sequencing. For instance, single cell skimming is possible since only 30 Mb per sample is needed. With low-cost enabled by sample multiplexing, genome skimming becomes ideal for epidemiological exposure monitoring for small magnitude global changes in DNA methylation in a population. Bioinformatic pipelines are maturing, and global methylation is expected to become a commonly reported metric for every type of nanopore run in the future.

### Biological and Technical Replication

To validate the level of precision necessary, I sequenced five biological replicates of three mouse tissues at skimming coverage levels (0.0025X to 0.016X). Detection range and accuracy are confirmed by repeated measures of the technical replicate control samples.

Biological replication is validated by highly consistent interindividual measurements among multiple tissues in the mouse biological replicates. Compared to other methods of quantifying DNA methylation, genome skimming is the most accurate and reproducible. While other methods exist to quantify global DNA methylation, none have the combination of ease of use, low-cost, low barrier to entry, high precision, and accuracy as nanopore genome skimming.

Currently the most common method for global DNA methylation uses a colorimetric antibody based commercial ELISA kit, which suffers from poor resolution with small magnitude differences and poor repeatability overall.^28^ Further, it only measures the methylated cytosine to total cytosine ratio, making its relation to cytosines in CpG context difficult to interpret.

Liquid-chromatography uses native DNA degraded to single bases, losing positional information and suffers from similar challenges as the ELISA method. A recent study using liquid chromatography across the tree of life assessed DNA modifications and reported large variation in biological replicates. By nature of the assay it also reports only methylated cytosine to total cytosine ratio and is difficult to extrapolate to cytosines in CpG context.^29^

Global methods relying upon the Qiagen Pyromark line of pyrosequencers were formerly popular such as the luminometric assay (LUMA) and the LINE1, *Alu*, and mouse IAP amplicon assays.^15,30^ The LUMA assay relies upon isoschizomer methylation-specific enzymatic digestion followed by pyrosequencer quantitation.^31^ The LINE1 and *Alu* assays rely upon bisulfite conversion and determine methylation solely within transposon context, obscuring changes outside transposon regions.^32^ Both methods utilize pyrosequencers which are phasing out of production.

### Vertebrate Methylation

I skimmed 40 different vertebrate species across fish, amphibians, reptiles, mammals, and primates, generating global DNA methylation values and transposon methylation. A study using WGBS in multiple vertebrate dermal fibroblasts showed global methylation that recapitulates my results nearly exactly^34^. For instance, their canine had 64.17% methylation to my 63.92%. Their human was 70.70% to my human 70.77%. Their rabbit had 68.26% to my rabbit 68.19%. Interestingly, their mapping statistics were rather poor, ranging from 73.5% to 79.3% while my long read skims mapped between 93.4% and 98.8% (mean 97.6%). This is a function of the short reads used in WGBS along with difficulty mapping due to bisulfite conversion.

The most comprehensive previous study of animal methylation assayed 580 species and matches the global patterns presented here^35^. However, this impressive catalog was created using RRBS and is subject to the typical challenges caused by bisulfite conversion, as well as enrichment for CpG dense regions and lack of accurate transposon identification due to short reads and loss of mapping ability.

I found that genome size directly correlates to methylation, explaining 44% of the variance. Clade specific methylation results replicate the phyloepigenetic tree built by Haghani et al. which was made with a custom 40k epigenetic microarray used on 176 species.^33^

### Transposons

Genome skimming is effective for transposon identification because repeats are present in many thousands of copies per nuclear genome. Transposon methylation is often used as a proxy for global methylation, however repeat copies are not truly identical and are divided by diagnostic mutations down to the subfamily and individual insertion level. Long reads allow precise identification of repeat type and often identify their exact genomic coordinates. Prior methods to determine DNA methylation of transposons included pyrosequencing, which amplifies a random mixture of transposons of a particular family and quantitates methylation at few specific CpG sites that are of unknown conservation in the bisulfite amplicon, leading to error and poor replication. Even at 0.001X coverage, equating to just 3 Mb of sequence, I was able to accurately determine *Alu* transposon methylation in the mouse genome, with less than 1% deviation between biological replicates. Genome skimming has been used in plants to identify and classify the extent of transposon content, but until now has not been used to assess DNA methylation of transposons.^12^

I found biologically meaningful results in the mouse transposome, showing that ERV_class II elements are substantially hypomethylated in mouse testes compared to other tissues. This is relevant because the most mutagenic transposon in the mouse genome, the IAP element, is an ERV_class II and its hypomethylation in testes suggests germ-line mobilization.^15,36^

### Summary

Genome skimming with nanopore sequencers is highly effective for global DNA methylation and transposon measurement. Multiplexing along with the ability to pause sequencing allows flexible experimental design and mitigates cost concerns. With improved base calling software and bioinformatic pipelines, global DNA methylation will likely be a basic quality metric for any type of nanopore run. Genome skimming for epigenetics stands to become a burgeoning field.

## Methods

### DNA Extraction

DNA from primate buffy coat samples was extracted Qiagen DNA mini kit (cat. 51306) by the Southwest National Primate Center (SNPRC) and treated with Qiagen RNAse A. DNA from mouse tissues and all vertebrate muscle samples were extracted using Zymo quick-DNA miniprep plus kit (cat. D4068) by the author. Vertebrate skeletal muscle tissues were sourced from commercial meat suppliers or donated by research labs. The human muscle sample was obtained commercially from BioChemed Services (Winchester, VA) and was de-identified prior to purchase. Mice in this study were of the agouti strain, wild type allele sharing at least 93% similarity with C57bl/6^37^. All were males 6-8 weeks of age. Mice in this study were maintained in accordance with the Guidelines for the Care and Use of Laboratory Animals (Institute of Laboratory Animal Resources, 1996) and were treated humanely and with regard for alleviation of suffering. The study protocol was approved by the University of Minnesota Institutional Animal Care and Use Committee (IACUC). Primate samples were acquired from the SNPRC from incidental blood collection during routine veterinary care and are under the aegis of SNPRC’s IACUC approval.

Control DNA was generated by treatment of mouse liver gDNA extracted via Zymo quick-DNA miniprep plus kit. For low methylated control, 100 ng of liver gDNA used as input to a Qiagen Repli-g Mini kit (cat. 150023) for whole genome amplification, following manufacturer’s protocol. Resulting DNA was cleaned using Axygen AxyPrep MAG PCR Clean-Up Kit (cat. MAG-PCR-CL-50) (Corning, Glendale, AZ). DNA was incubated with 2X volume of magnetic bead solution on a shaker for 5 minutes, washed twice with 70% EtOH, and eluted with 100 μl of elution buffer. Yield was 10 μg per reaction. Highly methylated control DNA was generated using 10 μg liver derived gDNA as input in separate reactions with a Zymo CpG Methylase kit (cat. E2010). Output DNA was cleaned with Axygen beads and yield was 70% of input. Control DNA was then used for library prep and sequencing like any other sample extract.

### Library preparation and run conditions

DNA from four primates was barcoded and prepped for sequencing using the Native Barcoding Kit SQK-NBD114.24 following manufacturer’s instructions from Oxford Nanopore Technologies (Oxford, UK). The marmoset sample was prepared with the EXP-NBD104 barcoding kit and the SQK-LSK109 ligation sequencing kit. Primate DNA was sequenced at the University of Wisconsin Biotechnology Center, Next Generation Sequencing Core on a PromethION 24 instrument. Two samples were multiplexed per PromethION flow cell.

All tissues from mouse and vertebrate muscle were library prepped using the Rapid Barcoding Kit SQK-RBK114.24 following manufacturer’s instructions by the author at the University of Minnesota. Mouse tissue DNA replicates along with control DNA were run in 24-sample multiplex along with technical replicates of five low and four high methylation individually barcoded controls from the same control reaction aliquots on a single MinION R10.4.1 flow cell for 24 hours. Vertebrate tissue DNA was run in 24 sample multiplex on a second MinION R10.4.1 flow cell for 24 hours. All MinION runs were performed in the author’s lab.

### Alignment and Methylation Assessment

Complete bioinformatic pipeline is available in the supplementary material and at https://faulk-lab.github.io/skimming/. Briefly, Guppy v6.3.8 was used to call bases for all species with the super accuracy model ‘dna_r10.4.1_e8.2_400bps_modbases_5mc_cg_sup’ that also calls 5’methycytosine in CpG context. Guppy generates mapped or unmapped bam files, depending on reference given. Here the reads were aligned post-hoc with minimap2, retaining modification flags by converting the bam files to fastq with samtools v1.16.1 and mapping with minimap2 to create a mapped bam file. Mapping efficiency was calculated using bamUtils v1.0.16. Next, the bam files were converted to bed format with modbam2bed 0.6.3. Summary methylation was calculated with an awk script available in the detailed supplementary methods. Read and genome summary statistics were calculated with seqkit. All bam files containing modified base calls are available at NCBI accession PRJNA927034. Differences in mouse tissue methylation were calculated with an ordinary 1-way ANOVA followed by Tukey’s multiple comparison test. Differences in low vs. high control methylation were calculated with an unpaired t-test. All mouse sequencing quality tests were compared with ordinary 1-way ANOVAs with Tukey’s multiple comparison test.

All software was run on a single computer running Ubuntu v22.04 on an AMD 7950 Ryzen 32 thread CPU with 128 Gb memory, 4 Tb SSD, and an Nvidia 3080 Ti GPU for accelerated base calling. All statistics and figures were created using GraphPad Prism v9.4.1.

### Vertebrate statistics

Vertebrate methylation was calculated similarly to mouse and primates, with alignment to the reference genome for each species downloaded from NCBI with accession numbers available as supplementary file 1. Genome size vs. methylation excluded axolotl as an outlier since its genome is an order of magnitude larger than any other vertebrate at 28 Gb. However, fitting with the pattern, it has the highest methylation percentage of any vertebrate sequenced.

### Transposon analysis

Mouse (mm39) and Chimpanzee (panTro6) repeatmasked tracks were downloaded from UCSC Genome Browser’s Table Browser function with the following parameters, “Table Browser -> Repeats -> Table “rmsk” -> All fields from selected table -> repeat-table.tsv”. The table was converted to bed format with a custom awk script. Bedtools was used to intersect the genome-wide methylation bed file generated with modbam2bed vs. the repeatmask track bed file. An R script was used to summarize the methylation values by repeat family and statistics applied were applied in GraphPad Prism. Mouse transposon family significance was calculated with a 2-way ANOVA followed by Tukey’s multiple comparison test. All scripts are available in supplementary file 2.

## Supplemental Data

Supplementary File 1 (Full data set)

Supplementary File 2 (Methods)

## Author contributions

CF conceived the study, performed the sequencing, analyzed the data, and wrote the manuscript. The author has read and approved the manuscript prior to submission.

