## Supplementary File 2 for "Genome Skimming with Nanopore Sequencing Precisely Determines Global and Transposon DNA Methylation in Vertebrates"

Genome Skimming


Code 

- Show All Code
- Hide All Code
- Download Rmd

### Genome Skimming

###### Chris Faulk

### Global DNA Methylation via Genome Skimming from Nanopore Long-read Data

Purpose: Most methods to characterize a global value for
5’methylcytosine are either expensive, error-prone, or rely upon
bisulfite conversion which causes measurement challenges. The nanopore
instrument can natively call 5’methylation along with other
modifications. Here I present a method of determining a single
percentage value of any genome’s 5’mCpG vs. unmodified CpG sites (i.e.
global methylation) by sequencing genomes at very low coverage (i.e.
skimming) at ranges from 0.1X to 0.0001X. Separately, I extracted high
copy transposon families and assessed their methylation
independently.

#### Coverage Depth Estimation

Genome skimming is defined as shallow sequencing down to 0.05X
coverage of a genome (0.05X). For a typical 3 Gb mammal genome this
equates to 150 Mb of reads. Here I show that for global DNA methylation,
even 0.01X coverage of a primate genome results in an error of less than
1% difference from the true value. The sampling was bootstrapped 10
times to determine error.

##### Subsampling Methods

There are approximately 29 million CpG sites in the human genome and
to get sub-percentile accuracy of their methylation, a measure of
methylation at 300,000 sites will suffice. Since the genome is
non-uniform with respect to CpG density and location, converting this
number into base coverage is not uniform. Here I chose to quantify
methylation on number of total bases sequenced since that is how a
sequencer counts reads and how the user determines how much is enough
DNA sequenced for skimming or assembly or other purposes. To estimate
whether a certain level of coverage will contain enough CpG sites for
accurate methylation, it was necessary to empirically sample the reads,
quantify methylation, and repeat the subsampling by bootstrapping to be
confident that a global value is correct.

Here I used the chimp genome as one of 4 primate genomes I sequenced
to high depth. The full dataset averages 77.8883% methylation over
11.12X coverage (33,921,056,788 bp) in the mapped bam file. Subsamples
were taken at 10X, 1X, 0.1X, 0.01X and 0.001X coverage and with 10
random subsets drawn, and methylation calculated for these 10 bootstrap
replicates.

- **10X** = 30504547470 bp = 10 \* (33921056788 /
  11.12). Therefore 0.899 \* total bases) = 10X. Sample 10X coverage 10
  times.
- Determine coverage.
  `mosdepth -t 32 --fast-mode rhesus.barcode02 chimp.modmapped.bam`
- Downsample the bam file with replicates.


```
for i in {1..10}; do samtools view -@ 32 -b --subsample 0.899 --subsample-seed $RANDOM total-chimp.modmapped.bam > bootstrap10X/total-chimp.modmapped.bam.10X.$i.bam ; samtools index -@ 32 bootstrap10X/total-chimp.modmapped.bam.10X.$i.bam ; done
```


- Convert bams to bed.


```
for i in bootstrap10X/*.bam; do modbam2bed -m 5mC -t 32 -e -p $i --aggregate --cpg panTro6.fa $i > $i.bed; done
```


- Use awk to average methylation.


```
for i in bootstrap10X/*.acc.bed ; do awk '$5>0 {can+=$12; mod+=$13} END{print 100*(mod/(can+mod))}' $i >> bootstrapped.10X.txt ; done
```


- **1X** = 30504547470 bp = 1 \* (33921056788 / 11.12).
  Therefore 0.090 \* total bases) = 1X. Sample 1X coverage 10 times.


```
for i in {1..10}; do samtools view -@ 32 -b --subsample 0.090 --subsample-seed $RANDOM total-chimp.modmapped.bam > bootstrap1X/total-chimp.modmapped.bam.1X.$i.bam ; samtools index -@ 32 bootstrap1X/total-chimp.modmapped.bam.1X.$i.bam ; done
```


- Repeat steps as in 10X for 1X .. 0.0001X

#### Biological and Technical Replication at Low Coverage

I used 5 biological replicates of 3 tissues from mouse to determine
intersample precision, i.e., reproducibility. I also included replicates
of high and low methylated control DNA.

##### Control Synthesis Methods

###### Low Methylation control DNA

This input was derived from mouse genomic DNA extracted from liver
tissue using a Zymo Quick DNA miniprep plus kit (cat. D4068). A small
amount of DNA was used as input to a Qiagen Repli-g whole genome
amplification kit (cat. 150023) according to instructions. The resulting
DNA amplified DNA is unmethylated, however the sample still contains a
small amount of the original mouse derived natively methylated DNA,
hence the name “low methylation control” rather than “0%”. The DNA was
then library prepped with ONT’s LSK-114 kit and sequenced on an R10.4.1
flow cell at 400 bps. Bases were called by Guppy v6.3.9 with the
`dna_r9.4.1_450bps_modbases_5mc_cg_sup.cfg` model.

###### High Methylation Control DNA

This sample was derived from mouse genomic DNA extracted from liver
tissue using a Zymo Quick DNA miniprep plus kit (D4068). The DNA was
subjected to CpG methylase treatment to induce high levels of 5’mCpG
using a Zymo CpG methylase kit (E2010) according to instructions. Due to
enzymatic limitations, the kit will not achieve perfect 100%
methylation, hence the name “high methylation control”. The DNA was then
library prepped with ONT’s LSK-114 kit and sequenced on an R10.4.1 flow
cell at 400 bps. Bases were called by Guppy v6.3.9 with the
`dna_r9.4.1_450bps_modbases_5mc_cg_sup.cfg` model.

##### Skimming Pipeline

Pipeline for analysis is as follows:

1. Optional. If you did not choose to call live during the run with SUP
   accuracy and DNA methylation detection turned on, or if you want to use
   a different model, you can re-basecall with guppy post-hoc using the
   original fast5 files. Guppy v6.3.8 was used here. NVIDIA GPU with CUDA
   cores is highly recommended to speed up base calling.


```
# Actual run command
/opt/ont/guppy/bin/guppy_basecaller -i /mnt/synology/ont-sequencing-run-storage/Skimming-methylation/mouse-skim-hip-cortex-testes-reps/no_sample/20230116_1229_MN34646_FAV16314_694e180d/fast5_pass/ -s . -c dna_r10.4.1_e8.2_400bps_modbases_5mc_cg_sup.cfg --bam_out -x cuda:all --recursive --trim_adapters --barcode_kits SQK-RBK114-24 --do_read_splitting

# Basic run command
opt/ont/guppy/bin/guppy_basecaller -i /path/to/fast5_pass/ -s /output/path -c dna_r9.4.1_450bps_modbases_5mc_cg_sup.cfg --bam_out -x cuda:0
```


2. Concatenate all the unmapped bam files containing modified base
   information.


```
samtools cat -o total.bam *input.bam
```


3. Map unmapped Bams to a reference. Map, convert and sort all in one
   command. The `-T Mm,Ml` flag carries over the modifications
   from bam to fastq. In newer bams the flags are
   `-T MM, ML`.  
   The `-y` flag copies input fastq commands to output.


```
# Convert unmapped bam to fastq, pipe output to minimap2, pipe output to samtools for sorting and indexing without writing intermediate files to disk. (Need sufficient RAM)

samtools fastq -T MM,ML total-chimp.bam | minimap2 -y -ax map-ont panTro6.fa - -t 32 | samtools view -u - | samtools sort -@ 32 -o total-chimp.modmapped.bam; samtools index total-chimp.modmapped.bam -@ 32
```


4. Convert to bed format. The `-d` flag limits depth to save
   memory. The aggregate flag adds together bases on either strand and
   creates and aggregate.bed file. The `-cpg` flag limits to cpg
   sites.


```
# Convert mapped bam file reads to .bed format to quantiate methylation by position.
modbam2bed -m 5mC -t 32 -e -p total-chimp.modmapped --aggregate --cpg panTro6.fa total-chimp.modmapped.bam > total-chimp.modmapped.bed
```


5. Determine methylation percentage by counting all canonical (5CpG)
   and modified (5mCpG) sites and dividing. Use the
   `mod-counts.cpg.acc.bed` file as it aggregates cytosine calls
   from both palindromic sides of the opposite strands.


```
# For every CpG dinucleotide, add up the count of canonical unmodified cytosines (column 12) and the 5'methylated cytosines (column 13) at every mapped position. Divide the number of modified cytosines by the total number of cytosines.

# Note that modbam2bed produces a stranded and a combined "acc" file. The combined file adds together the opposite strand cytosine counts together since they represent the same palindromic cytosine. This increases power.

awk '$5>0 {can+=$12; mod+=$13} END{print 100*(mod/(can+mod))}' total-chimp.modmapped.cpg.acc.bed
```

### Vertebrate Genomes

DNA from species representing the full vertebrate radiation were
collected through commercial suppliers and gifts from collaborators. All
samples were derived from muscle tissue and extracted using Zymo Quick
DNA Plus miniprep kits. Genomes were downloaded from Genbank with the
latest reference available for each species.

1. Merge and copy the bams for each barcode.


```
samtools cat -o total.bam *input.bam
```


2. Map to genome.


```
samtools fastq -T MM,ML total-chimp.bam | minimap2 -y -ax map-ont reference.fa - -t 32 | samtools view -u - | samtools sort -@ 32 -o animal.modmapped.bam; samtools index animal.modmapped.bam -@ 32
```


1. Use `--split-prefix` if the genome is too large for
   minimap2.
2. Convert to bed file.


```
modbam2bed -m 5mC -t 32 -e -p reference.modmapped --aggregate --cpg reference.fa animal.modmapped.bam > animal.modmapped.bed
```


4. Calculate Methylation.


```
awk '$5>0 {can+=$12; mod+=$13} END{print 100*(mod/(can+mod))}' animal.modmapped.cpg.acc.bed
```


5. Calculate mapping efficiency.
6. Basic stats
   `bam stats --in animal.modmapped.bam --basic`
7. Average quality
   `samtools stat animal.modmapped.bam | grep "average quality"`
8. Copy to Excel and Graphpad for graphics.

### Transposon Analysis

#### Primate LINE1 and Alu elements

These 2 elements alone account for ∼30% of the genome sequence and
are the most abundant transposable elements in humans (Lander et
al.)

#### All Repeats Genome-wide

0.8604273% Average methylation across all repeat types

#### Transposon Pipeline

###### Chimpanzee LINE1 and Alus Specifically

1. Table Browser -> Repeats -> Table “rmsk” -> All fields from
   selected table -> repeat-table.tsv
2. Get all L1’s from Repeats tsv and print into bed format.


```
awk '$13=="L1" {print $6 "\t" $7 "\t"$8 "\t"$13}' panTro6-repeat-table.tsv > repeatL1.bed
```


3. Intersect all CpG sites that fall within L1 coordinates and average
   their methylation


```
bedtools intersect -a total-chimp.modmapped.cpg.acc.bed -b repeatL1.bed > repeatL1.cpgintersect.bed
awk '$5>0 {can+=$12; mod+=$13} END{print 100*(mod/(can+mod))}' repeatL1.cpgintersect.bed
```


1. 84.7069% L1 methylation in the 11.12X coverage chimp
   genome.
2. 92.4767% Alu in the chimp genome.
3. Downsample the reads to 0.001X (3 Mb) and plot 10 bootstrap
   replicates.
4. Use the pre-made downsampled cpg.acc.bed files in the bootstrap
   directories
5. Make intersection files for each bootstrap.


```
```bash
for i in bootstrap10X/*.cpg.acc.bed ; do bedtools intersect -a $i -b repeatL1.bed > $i.cpgintersect.L1.bed ; done
```
```


3. Use awk to print the methylation to a file.


```
```bash
for i in bootstrap10X/*.cpgintersect.bed ; do awk '$5>0 {can+=$12; mod+=$13} END{print 100*(mod/(can+mod))}' $i >> L1bootstrapped.10X.txt; done
```
```


4. Repeat for each read level.
5. Repeat for Alus.

###### Chimpazee All Repeats

1. Chimp. Make an intersection of all repeats and cpg sites with -wa
   and -wb to combine beds.


```
bedtools intersect -a total-chimp.modmapped.cpg.acc.bed -b repeatAll.bed -wa -wb > repeatAll.cpgintersect.bed
```


2. Mouse. Repeat as with Chimp, but use all 5 mouse hippocampus samples
   that share common rows for methylation data.

##### Mouse Repeats

###### Mouse All Repeats

1. Download all mm39 repeats from USCS Table Browser.
2. Convert to bed format.


```
awk '{print $6 "\t" $7 "\t"$8 "\t"$13}' mm39-repeat-table.tsv > mm39-repeat-table.bed
# Remove the header line manually.
```


3. Make an intersection of all repeats and cpg sites with -wa and -wb
   to combine beds. Loop for all files.


```
for i in *.cpg.acc.bed ; do bedtools intersect -a $i -b mm39-repeat-table.bed -wa -wb > $i.cpgintersect.bed ; done
```


4. Use R program `repeats-rmd.Rmd` (copied below) to
   summarize methylation on a per-repeat basis and copy results into
   Graphpad for analysis and visualization.


```
```r
### Import necessary libraries
library(tidyverse)

### Read in tsv filea
data <- read_tsv("mouse-hip.modmapped.cpg.acc.bed.cpgintersect.bed", col_names=FALSE)

### Calculate sum of column 13 divided by the sum of column 12 plus column 13
data$result <- data$X13 / (data$X12 + data$X13)
data <- data %>% drop_na()
data$result

### Separate out by factors in column 18
data_by_factor <- data %>% group_by(X18) %>% summarize(result = mean(result))
data_by_factor
write.table(data_by_factor , file = "mouse-hip.modmapped.cpg.acc.bed.cpgintersect.bed.databyfactor.tsv")

### Make a bar plot of the result
ggplot(data_by_factor, aes(x = X18, y = result)) + geom_bar(stat = "identity")  + 
  theme(axis.text.x = element_text(angle = 90, vjust = 0.5))
```
```
